# A whole virion vaccine for COVID-19 produced via a novel inactivation method: results from animal challenge model studies

**DOI:** 10.1101/2020.11.13.381335

**Authors:** Izabela K Ragan, Lindsay M Hartson, Taru S Dutt, Andres Obregon-Henao, Rachel M Maison, Paul Gordy, Amy Fox, Burton R Karger, Shaun T Cross, Marylee L Kapuscinski, Sarah K Cooper, Brendan K Podell, Mark D Stenglein, Richard A Bowen, Marcela Henao-Tamayo, Raymond P Goodrich

## Abstract

The COVID-19 pandemic has generated intense interest in the rapid development and evaluation of vaccine candidates for this disease and other emerging diseases. Several novel methods for preparing vaccine candidates are currently undergoing clinical evaluation in response to the urgent need to prevent the spread of COVID-19. In many cases, these methods rely on new approaches for vaccine production and immune stimulation. We report on the use of a novel method (SolaVAX™) for production of an inactivated vaccine candidate and the testing of that candidate in a hamster animal model for its ability to prevent infection upon challenge with SARS-CoV-2 virus. The studies employed in this work included an evaluation of the levels of neutralizing antibody produced post-vaccination, levels of specific antibody sub-types to RBD and spike protein that were generated, evaluation of viral shedding post-challenge, flow cytometric and single cell sequencing data on cellular fractions and histopathological evaluation of tissues post-challenge. The results from this study provide insight into the immunological responses occurring as a result of vaccination with the proposed vaccine candidate and the impact that adjuvant formulations, specifically developed to promote Th1 type immune responses, have on vaccine efficacy and protection against infection following challenge with live SARS-CoV-2. This data may have utility in the development of effective vaccine candidates broadly. Furthermore, the results suggest that preparation of a whole virion vaccine for COVID-19 using this specific photochemical method may have utility in the preparation of one such vaccine candidate.

**Author Summary:** We have developed a vaccine for COVID-19 which is prepared by a novel method for inactivation of a whole virion particle and tested it in a hamster animal model for its ability to protect against SARS-CoV-2 infection.

## Introduction

Coronavirus disease 2019 (COVID-19), caused by the virus SARS-CoV-2, continues to spread globally leading to significant impacts on public health [1]. As of August 2020, over 29 million total cases have been confirmed and over 900,000 deaths have been reported globally [2]. SARS-CoV-2, or Severe Acute Respiratory Syndrome coronavirus 2, causes primarily respiratory infections in humans and is related to other coronaviruses like Middle East Respiratory Syndrome Coronavirus (MERS-CoV) and SARS-CoV [3]. Not only has SARS-CoV-2 caused a public health emergency on a global scale but it continues to have major social, cultural, and economic impacts. Vaccination is the most effective countermeasure for mitigating pandemics and has proven effective against viral pathogens such as smallpox and polio.

The emergence of SARS-CoV-2 in human populations spurred the scientific community to development methods to produce vaccines against COVID-19 [4]. Many development paths were initiated using methods that have been applied historically to the production of vaccine candidates as well as new approaches aimed at delivering candidates in a rapid and efficient manner [5]. These methods have included the use of RNA and DNA vaccines, subunit vaccines, attenuated vaccines, as well as vectored vaccines utilizing virus-like particles (VLP), adenovirus or bacterial host constructs. Inactivated vaccines have been a mainstay of vaccinology for decades. Even today, examples of inactivated vaccines include constructs for influenza, cholera, bubonic plague and polio [6]. Each method of vaccine production has merits and limitations. These limitations range from issues in production at scale, cost to produce, stability and delivery process and the potential for side effects.

The use of these vaccine production methods is predicated on the ability to achieve inactivation of the pathogen’s ability to replicate while preserving the antigenic protein structural potency and integrity [7]. Methods for producing such candidates require the use of chemicals such as beta-propiolactone, ethylenimine, formalin as well as the use of physical inactivation methods such as gamma irradiation, high energy UV light (UVC) and heat inactivation. These methods work primarily by chemically modifying protein and nucleic acid structures of pathogens through covalent modification, crosslinking, oxidation and structural alteration, rendering the treated agents unable to infect target cells or replicate in vivo. The key to success with such approaches resides in the ability to balance the alterations required for inactivation with the preservation of antigen epitopes similar to the native agent that are required to stimulate immune response when administered to a host. A method which can successfully prevent pathogen replication without inducing alterations to antigen targets is ideal. Candidates produced by such a method would possess the native protein structures that are as close to natural structures in the intact pathogen as possible without the ability to induce disease.

We report a novel method (SolaVAX™) to produce a candidate vaccine for COVID-19. This method employs the use of a photochemical (riboflavin or vitamin B2) in combination with UV light in the UVA and UVB wavelength regions to carry out specific nucleic acid alterations through electron transfer chemistry-based processes [8]. The method was originally developed for the treatment of blood products to prevent transfusion transmitted diseases [9] and has been in routine clinical use for prevention of transfusion transmitted viral, bacterial and parasitic diseases since 2008 [10]. The process utilizes a well-established and demonstrated capability of riboflavin and UV light to modify nucleic acid structure primarily through modification of guanine bases in a non-oxygen dependent process utilizing the natural electron donor-acceptor chemistry associated with guanine and riboflavin, respectively [11].

The specificity of the riboflavin photochemistry used in this process avoids the alkylation, crosslinking and covalent modifications that are associated with other chemical and photochemical mechanisms of pathogen inactivation [12]. The process allows for retention of plasma and cell-bound protein structure post treatment to an extent that such products may still be efficacious in functional utilization for transfusion support of patients [13,14]. Unlike the standard chemical agents such as beta-propiolactone and ethyleneimine derivatives that are routinely used for inactivated vaccine production, the photochemical used in this approach (riboflavin) has a well-established safety toxicology profile, is non-mutagenic and non-carcinogenic and poses little to no toxicity or disposal risk to facility personnel or the environment. This safety profile has been documented extensively in pre-clinical and clinical programs in human subjects [15].

For this work, we hypothesized that the use of these methods for production of an inactivated virus of SARS-CoV-2 could have several advantages. These included the ability to utilize existing equipment, reagents and disposables that are in routine use for treatment of blood products to produce an inactivated vaccine preparation when using purified viral stocks of the target virus, e.g., SARS-CoV-2 in this particular application [16]. We hypothesized that the selectivity of the chemistry applied in this process would generate a vaccine candidate that was fully attenuated with regard to replication capabilities while maintaining viral protein structural integrity as close to the native virus as possible. We further hypothesized that such a candidate produced by this method would be able to induce a potent immune response with relatively low doses of immunogen and thus provide protective immunity against live virus challenge.

Our approach to evaluate this candidate utilized both *in vitro* analysis and animal models (Syrian Golden Hamsters), developed in house and shown to be susceptible to disease in the upper and lower respiratory tract following exposure to live SARS-CoV-2 virus administered intranasally. Animals exposed in this way were demonstrated to produce and shed virus extensively at days 3-7 post-exposure. Our methods for evaluation of an effective vaccine candidate in this model included measurement of neutralizing antibodies with a Plaque Reduction Neutralization Test (PRNT), assessment of tissue viremia via plaque assays for live virus, flow cytometric analysis of leukocyte subpopulations, single cell mRNA sequencing analysis to assess host transcriptional responses and histopathology of respiratory system post-challenge with live virus via the intranasal route.

As part of these studies, we also utilized adjuvant formulations intended to drive immune response to vaccines predominantly via a Th1 immune pathway [17,18]. Our motivation for employing this approach was generated by previous observations of antibody dependent enhancement (ADE) and resulting immunopathology in animal models where Th2 type immune stimulation predominated. We monitored immune response via cellular and humoral pathways in adjuvanted and non-adjuvanted formulations. We also monitored lung histopathology for evidence indicative of immunopathology induced by the virus or subsequent challenge post-vaccination.

## Results

### Sequence validation of SARS-CoV-2 isolate

We used shotgun RNA sequencing to confirm the identity of the SARS-CoV-2 isolate used in hamster challenge material and to characterize mutations that may have arisen during cell culture passage [19,20]. The isolate contained 5 consensus-changing mutations relative to the USA-WA1 patient-derived sequence (GenBank accession MN985325.1) [21] from which it was derived (Table S1). These included a G to A substitution at position 23616 of the genome that results in an arginine to histidine substitution at position 685 within the polybasic cleavage site of the spike protein.

### Assessment of RNA damage following UV treatment

We used sequencing to detect and quantify RNA damage in photoinactivated virus preparations. Damaged RNA bases can be misrecognized during reverse transcription, leading to characteristic mutations in cDNA sequences [22]. We detected evidence consistent with oxidative damage to G bases in viral RNA in the form of elevated frequencies of G to U and G to C mismatches (Fig. 1B). These lesions would result from the misincorporation of an A or a G opposite an oxidized G during reverse transcription. The frequency of G to U mutations was 2.3x higher in photoinactivated RNA than in untreated RNA (0.0021 vs. 0.0009), and this ratio was 1.8x for G to C mutations (0.0012 vs 0.0007; Fig. 1C). Given the combined mutation frequency of ∼0.0033 and 5,863 G bases in the SARS-CoV-2 USA-WA1 genome, an average of 19.6 G bases will be damaged per genome. Using these parameters to estimate the number of damaged G bases per genome with a Poisson distribution estimated that 1 genome per 3.0×10^9^ genomes will have no damaged bases (Fig. 1D).

**Fig. 1.**
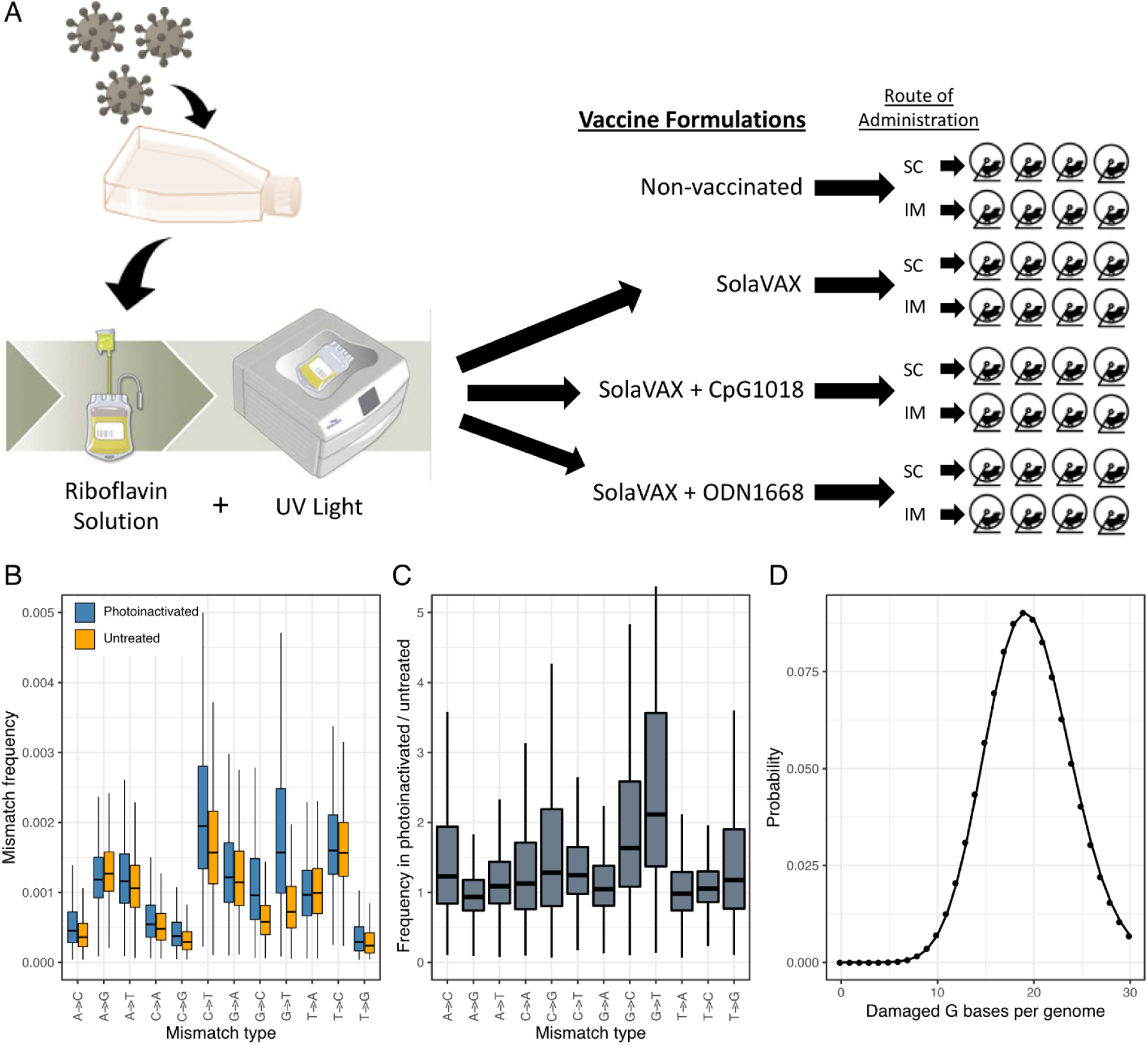
Production of SolaVAX vaccine and sequence-based detection of RNA damage in photoinactivated virus preparations. **(A)** SARS-CoV-2 virus (isolate USA-WA1/2020) was propagated in Vero E6 cells. The virus was then inactivated using the Mirasol PRT System by adding a riboflavin solution to the virus stock and exposing the solution to UV light. The inactivated virus was concentrated and prepared with or without adjuvant (CpG 1018, ODN1668). Hamsters were immunized with various SolaVAX vaccine formulations either subcutaneously (SC) or intramuscularly (IM) in groups of four animals. **(B)** The frequencies of the indicated mismatches in SARS-CoV-2 mapping reads in datasets from untreated or photoinactivated virus preparations. Mismatches are relative to the SARS-CoV-2 positive sense RNA sequence. **(C)** The ratio of mismatch frequencies in inactivated and untreated datasets, normalized to the frequency of bases in the reference sequence. Boxplots represent distributions of values across all sites in the genome. **(D)** A Poisson distribution estimating the probability of a SARS-CoV-2 genome containing the indicated number of damaged G bases, assuming 5863 Gs per genome and a combined mismatch frequency of 0.0033 for G to C and G to U mutations.

### Hamster model

Hamsters were divided into 4 treatment groups in which Control hamsters received no vaccination, a second group (SvX) received inactivated vaccine (SolaVAX) with no adjuvant, the third group (CpG) received inactivated vaccine with CpG 1018 adjuvant, and the fourth group (ODN) received inactivated vaccine with ODN1668 adjuvant. Within each group hamsters were divided in 2 subgroups that were vaccinated either by subcutaneous (SC) or by intramuscular (IM) injection (Fig.1A). None of the hamsters showed any clinical adverse reactions to the vaccination. Moreover, after viral challenge all hamsters were clinically normal. From the time of viral challenge to necropsy a body weight loss of 4-7.2% was observed in all groups including the Controls.

### Virus titration

Hamsters were challenged with 10^5^ plaque forming units (pfu) of live SARS-CoV-2 intranasally. Then oral-pharyngeal swabs were taken 1-3 days post-infection (dpi) to monitor viral replication. At 1 dpi infectious virus was detected in all groups (Fig. 2A-B). At 2 dpi viral replication begins to decline especially in the vaccinated groups. And by 3 dpi, viral replication was detected only in the Control group (SC and IM) and the ODN cohort (SC only). This shows that vaccination in hamsters reduced the amount of viral replication in the oropharynx after SARS-CoV-2 infection.

**Fig. 2.**
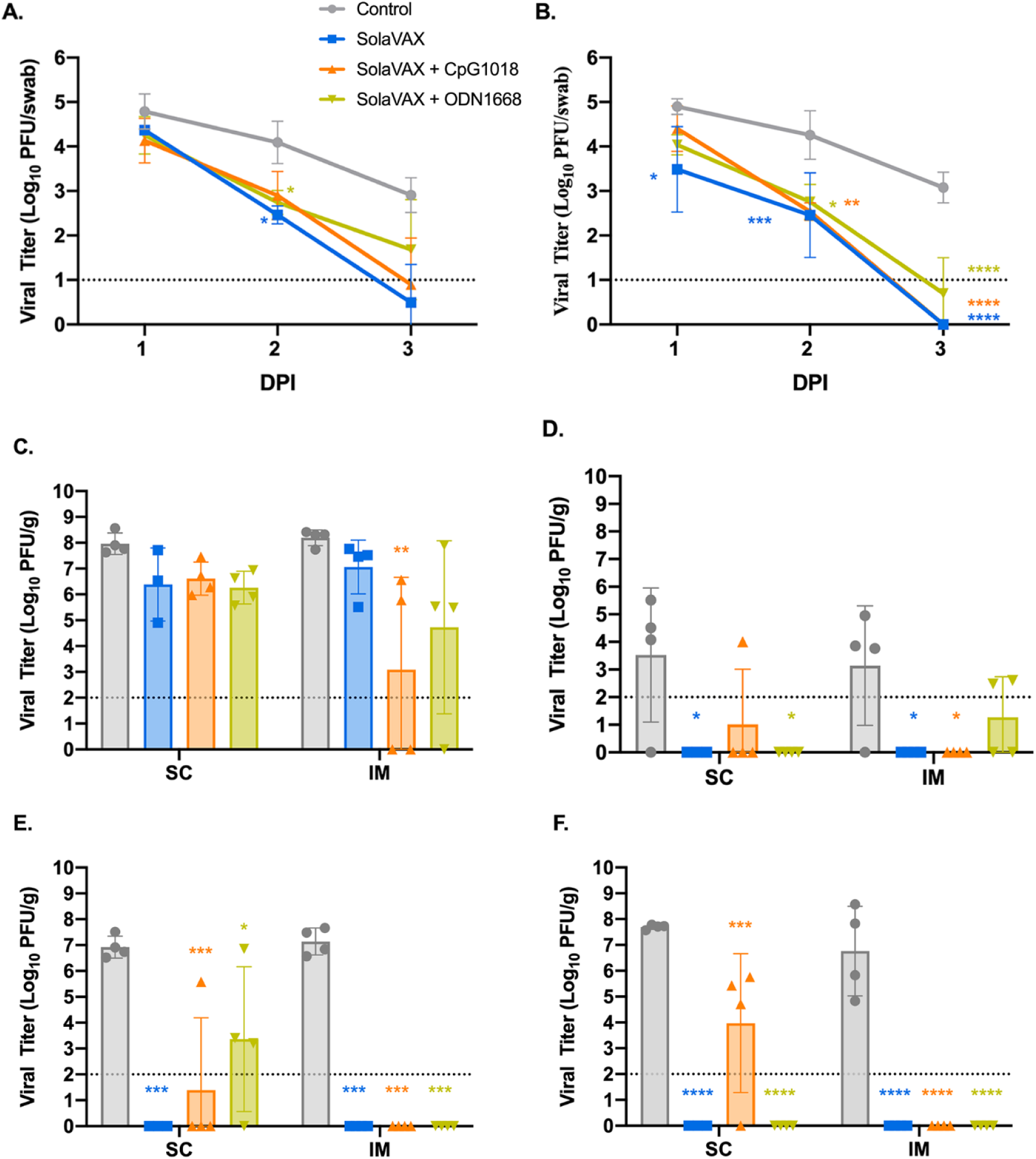
Viral loads from oropharyngeal swab and respiratory tract tissues after challenge with live virus. Oropharyngeal swabs were taken from all hamsters on 1, 2, and 3 days post infection (DPI). Viral titers of swabs collected from hamsters vaccinated via SC (**A**) and via IM (**B**) were determined by plaque assay. The presence of infectious virus was also determined in turbinates **(C)**, trachea **(D)**, right cranial lung lobe **(E)**, and right caudal lung **(F)** of each hamster three days after live virus challenge. Data points represent group mean +/- SD. SC, subcutaneous vaccination. IM, intramuscular vaccination. Asterisks above bars indicate statistically significant difference in viral titers between Control and vaccine group (**** =p<0.0001, *** =p<0.001, **=p<0.01, * =p<0.05). Limit of detection denoted as horizontal dotted line.

In addition to oral-pharyngeal swabs, necropsies were performed and tissues collected at 3 dpi to determine vaccine efficacy after live virus challenge. These tissues were specific to respiratory tract and included the right cranial lung lobe, right caudal lung lobe, trachea, and nasal turbinates Beginning with nasal turbinates (Fig. 2C), these tissue samples revealed high viral titers for all hamsters regardless of vaccination status. This was expected due to the route of live virus inoculation, yet the Control group had higher viral titers compared to the vaccinated groups. Moreover, there was a significant decrease in viral titers in the CpG group within the IM subgroup demonstrating that the IM injection of SolaVAX+CpG 1018 offered the best protection against viral replication in nasal turbinates.

Trachea was also evaluated to see if vaccination would protect the lower airway against SARS-CoV-2 infections (Fig. 2D). The viral titers are less than what was observed in the turbinates and support what has been seen in previous experimental hamster infections with SARS-CoV-2. Groups SvX and ODN showed a significant reduction in viral replication by SC administration while groups SvX and CpG showed a significant reduction by IM administration when compared to their respective Controls. In summary, SolaVAX+ CpG 1018 administered IM is effective in protecting against viral replication in both the nasal turbinates and trachea of hamsters.

Cranial and caudal lung lobes were collected to evaluate protection against SARS-CoV-2 in multiple lung lobes. Previous experimental infections with SARS-CoV-2 in hamsters revealed the viral load between the two lobes are usually similar, as was observed in the Control groups (Fig. 2E-F). However, the cranial lobe is commonly affected first before the caudal lobe. Therefore, it is of interest to evaluate the cranial lung lobe for vaccine efficacy against SARS-CoV-2 early in disease progression. Within the SC subgroup, all hamsters except for one hamster in the CpG cohort and three hamsters in the ODN cohort had no detectable virus (Fig. 2E). Within the IM subgroup, no infectious virus was detected in any of the vaccinated hamsters. Therefore, vaccination appeared to have reduced viral replication in the cranial lung lobes compared to non-vaccinated hamsters.

Lastly, the caudal lung was evaluated for the presence of infectious virus. As with the cranial lung, viral replication was detected in all hamsters in the Control group (Fig. 2F). Within the SC subgroup, only three hamsters within the CpG cohort In the IM subgroups, no viral replication was detected in any of the vaccinated groups. As seen with the cranial lung, all the vaccinated groups appeared to have reduced viral replication in the caudal lung lobes compared to non-vaccinated hamsters.

## Serology

### Plaque Reduction Neutralization Test

A plaque reduction neutralization test (PRNT) with a 90% cutoff was performed to measure neutralizing antibodies against SARS-CoV-2 after vaccination. Neutralizing antibodies were measured 21 days after the initial vaccination and then at 42 days after initial vaccination (21 days after booster vaccination). All hamsters were seronegative against SARS-CoV-2 prior to vaccination. As expected, hamsters in the Control group did not develop a detectable neutralizing antibody response against SARS-CoV-2 (Fig. 3A and B). In contrast, all but one hamster (CpG, SC) developed antibody titers ranging from 1:10-1:160 after the first vaccination. Moreover, there was an increase of antibody response in all but one of the vaccinated hamsters ranging from 1:40-1:1280. Two SvX hamsters (IM) had a detectable titer of 1:80 and 1:160 after first vaccination but no detectable titer after booster vaccination. Booster vaccination in general increased the titer of neutralizing antibodies prior to virus challenge. In comparing all vaccinated groups, CpG (IM) had the highest mean titer after both the prime and the booster vaccination.

**Fig. 3.**
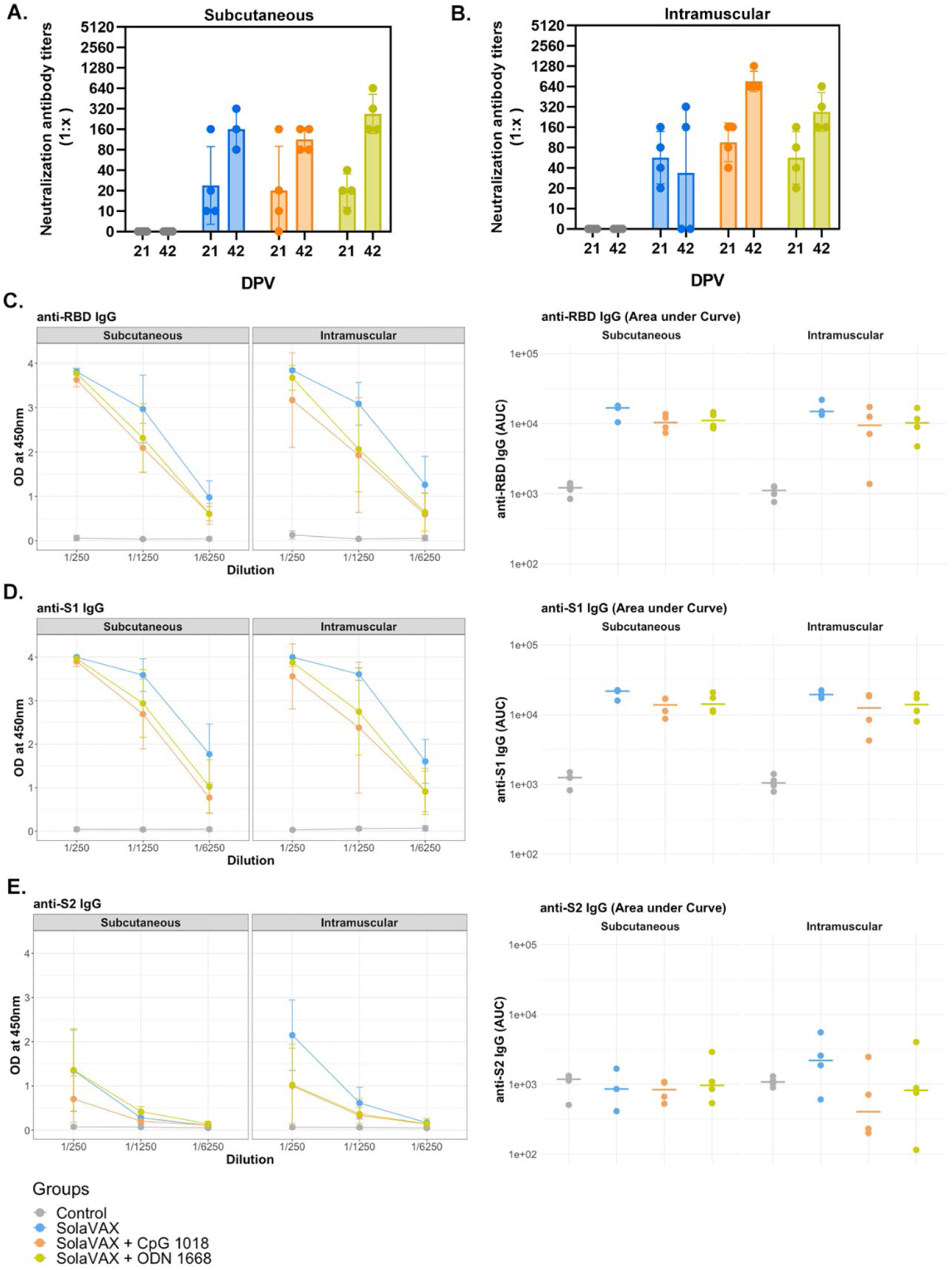
The detection of neutralizing antibodies in hamsters by PRNT90 after vaccination (A and B), and serum reactivity to spike protein’s RBD, S1 and S2 region (C-E). A plaque reduction neutralization test with a cutoff of 90% was used to determine neutralizing antibody production after 21and 42 days post vaccination (DPV) for both SC **(A)** and IM **(B)** routes of vaccine administration. The prime vaccination was given at 0 DPV and a booster vaccination given at 21 DPV. Data points represent group mean +/- SD. Results of ELISAs measuring serum reactivity to RBD **(C)** and S1 **(D)** and S2 **(E)** protein. Graphs on the left panel shows optical density (OD) at 450 nm (y-axis) vs serum dilutions (x-axis). Values represent mean +/- SD, n=4. Right panel shows area under the curve (AUC) calculated for each dilution for individual hamsters.

### ELISA

Enzyme-linked Immunosorbent Assay (ELISA) was performed on serum samples from Control and vaccinated hamsters to test for the presence of IgG against the SARS-CoV-2 spike protein’s receptor-binding domain (RBD) and spike protein regions S1 and S2 (Fig. 3C-E). Pooled serum from naïve hamsters was used as a negative control. A strong IgG response against the three viral proteins was detected in infected hamsters previously vaccinated with SolaVAX. In contrast, IgG levels against viral proteins were below the detection limit in hamsters in the Control group. The following trend was observed for all vaccinated hamsters regardless of vaccination route: a) Titers against the S1 protein and RBD subunit were higher than against the S2 protein; b) IgG levels were greater in infected hamsters vaccinated with SvX, followed by ODN and CpG groups. Panel B shows values for individual hamsters as an area under the curve. This suggests that neutralizing antibodies alone are not solely responsible for the enhanced protection provided by CpG group.

### Histopathology

Hematoxylin and eosin (H&E) stained slides, including sections of lung, trachea, heart and spleen, were reviewed for histopathological changes due to SARS-CoV-2 infection and alleviation of pathology through vaccination (Fig. 4). No significant pathology was identified in heart or spleen tissue. Control hamsters infected with SARS-CoV-2 demonstrated the most severe pulmonary pathology. Histopathological features of SARS-CoV-2 infection in this group included a strong predilection for larger airways including hilar bronchi and trachea. Bronchi and trachea contained lymphocytic inflammation infiltrating the mucosal epithelium and submucosa in seven of eight Control hamsters, accompanied by neutrophil dominated inflammation disrupting the epithelial surface or completely filling the airway lumen present in five of eight Control hamsters. Control hamsters also developed the most severe alveolar pathology. Alveolar walls were expanded by mononuclear inflammatory cell infiltrates, which limited alveolar air space, and in regionally extensive areas of the lung, led to consolidating interstitial pneumonia with complete effacement of normal alveolar structures. Inflammatory processes in the alveolar spaces were uniformly cell-mediated and lacked evidence of vasoactive inflammation including an absence of edema fluid and fibrin.

**Fig. 4.**
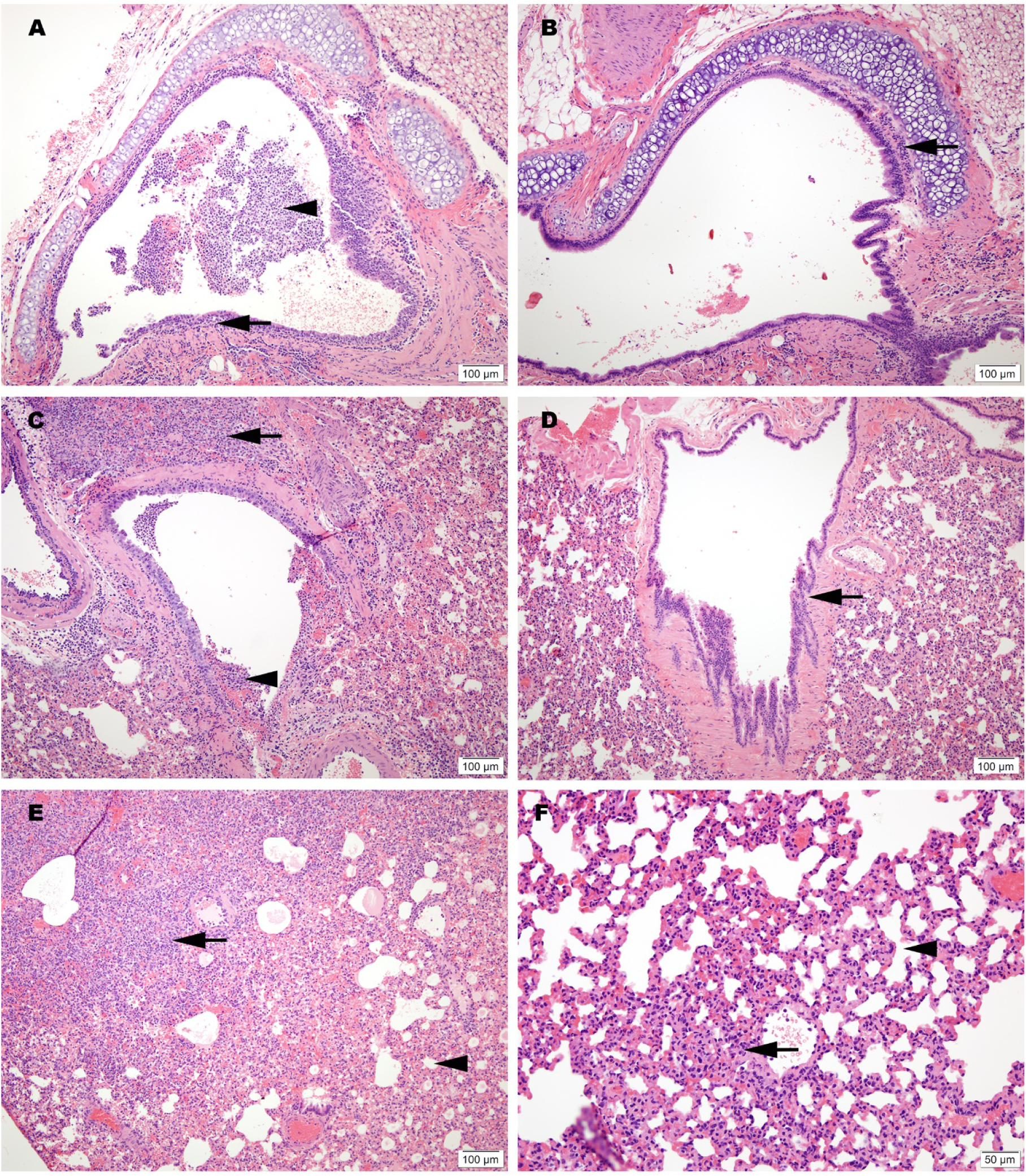
Representative histology of differences between unimmunized SARS-CoV-2 infected controls (A, C and E) and infected hamsters vaccinated with SolaVAX prepared SARS-CoV-2 virus and CpG 1018 adjuvant (B, D and F). **(A)** Trachea with dense submucosal lymphocytic and neutrophilic inflammation infiltrating mucosal epithelium (arrow) and accumulation of neutrophils within the tracheal lumen (arrowhead). **(B)** Trachea with mild submucosal lymphocytic inflammation. **(C)** Large bronchus with dense lymphocytic and suppurative inflammation in the interstitium (arrow) and accumulation of neutrophils in the lumen with loss of mucocal epithelium (arrowhead). **(D)** Large bronchus minimally affected by inflammation (arrow). **(E)** Effacement of lung alveolar tissue by consolidating interstitial pneumonia (arrow) and overall decrease in alveolar air space (arrowhead). **(F)** Interstitial pneumonia increasing alveolar wall thickness (arrowhead) without compromising alveolar air space (arrowhead).

Among vaccinated hamsters, those in the CpG (IM) group were the best protected from viral-induced pathology. Hamsters immunized with this formulation had improved air space capacity, a lack of consolidating inflammation, and bronchi or trachea with mild inflammatory changes or essentially normal morphology. ODN hamsters were also protected, but to a lesser extent than the CpG group. Notably, however, the SvX group offered a level of protection from severe pulmonary pathology compared to the Controls, and while not achieving statistical significance, this was observed primarily in hamsters vaccinated by SC route (Fig. S1).

### Immune Responses

The immunological response elicited upon SARS-CoV-2 infection of control or vaccinated animals was evaluated by flow cytometry analysis of leukocytes obtained from lungs, spleen and blood. Populations classified as having statistically significant differences in the total numbers of each cell type present in the lung/spleen or blood between groups are shown in Figure 5. In the lungs, the SC Control group had significantly more cells expressing inflammatory markers (IL-6) and (IL-6 and CXCR4) than any of the other subcutaneously vaccinated groups. The CpG SC group also had significantly fewer cells associated with a Th2 response (CD8+ IL-6+ GATA3+ CD4-CXCR3-CXCR4-IL-4-IL-10-Tbet-IFN-γ-TNF-α-) compared to the SvX vaccinated group. These cells may be involved in the induction of isotype switching in the host as a result of the increased infection in the Control group. The vaccine appears to shift the immune response away from an anti-inflammatory Th2 response. Furthermore, in the blood, the SolaVAX-vaccinated IM groups with adjuvants had significantly lower numbers of CD8+ CXCR4+ CD4-CXCR3-IL-6-Tbet-IFN-γ-IL-4-IL-10-GATA-TNF-α-cells compared to the Control group. These cells may promote inflammatory cytokine expression and cell chemotaxis through the MAPK pathway. In the spleen, the Control group had significantly higher numbers of proinflammatory cells expressing CXCR4 + IL-6+ Tbet+ IFN-γ + CD4-CD8-CXCR3-IL-4-IL-10-GATA-TNF-α-cells than the vaccinated groups. Non-significant populations for all organs are shown in Figure S2.

**Fig. 5.**
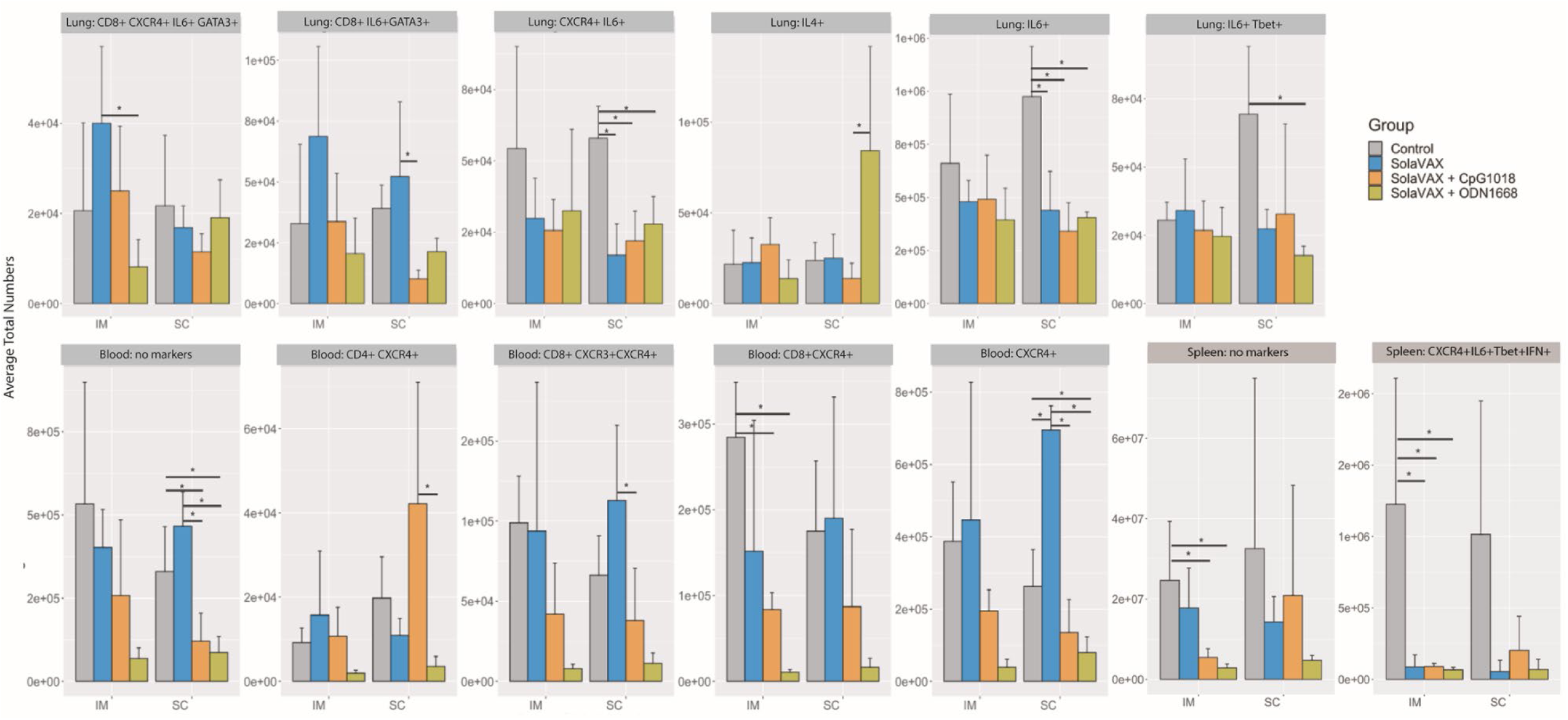
Statistically significant flow cytometry populations within intramuscular and subcutaneously vaccinated groups. The bar plots show the statistically significant populations identified through cyto-feature engineering for each organ. The x-axis shows the eight groups studied. The average total numbers of cells for each group were calculated and shown. The population names listed at the top of each small plot indicates the flow cytometry markers that are positive in the population. Note that these populations are negative for all of the other markers in the panel.

Flow cytometry analysis of the Syrian hamster immunological response was limited by the paucity of reagents available for this animal model. Thus, to get a better understanding of the immune response and antiviral or pathogenic mechanism(s) elicited by the different SolaVAX formulations, single cell transcriptomics (scRNAseq) analysis was performed on cells obtained from lungs of vaccinated or unvaccinated Syrian hamsters exposed to SARS-CoV-2-infection. Transcripts were detected from an average of 750 different genes with approximately 20,000 reads/cell (Fig. S5). Using an unsupervised cluster detection algorithm (SEURAT) at low resolution, four cellular clusters were identified by the lineage-defining genes CD3D (T cells), CD86 (Myeloid cells), MARCO (Myeloid cells), SFTPC (Epithelial cells), and CD79B (B cells) (Fig. 6 A-C). All the genes used to identify cell types are presented in supplementary table S2. Consistent with the histopathological analysis (Fig. 4), the myeloid population was increased in non-vaccinated hamsters. Similarly, there was a higher relative abundance of T cells in lungs of CpG hamsters, in agreement with the flow cytometry results. Epithelial cells were more abundant in all SvX-vaccinated groups, especially in the CpG group, consistent with increased abundance of inflammatory cells in non-vaccinated hamsters (Fig. 6).

**Fig. 6.**
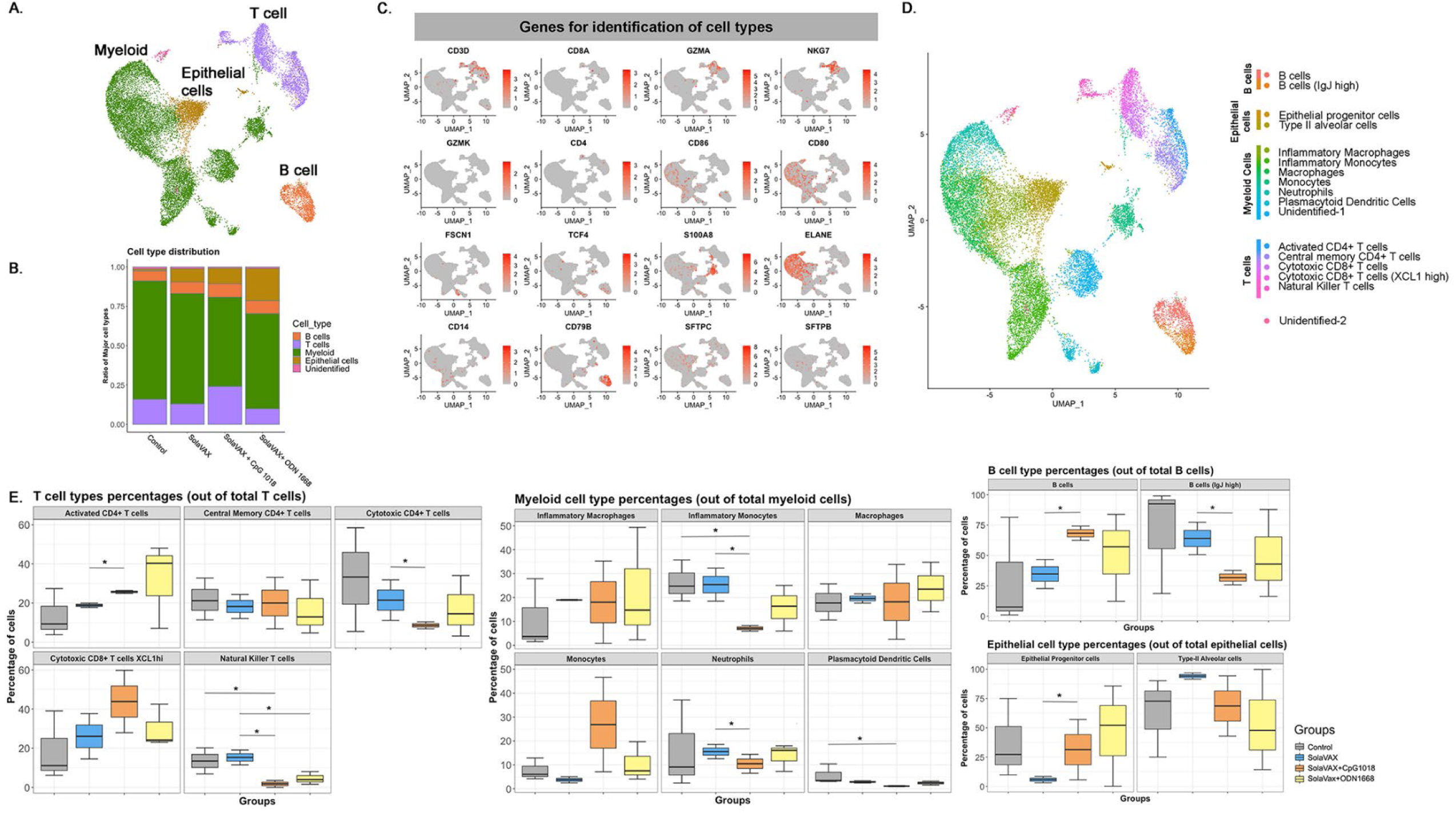
Single-cell transcriptomes of lungs from non-vaccinated and SolaVAX vaccinated hamsters. **(A)** merged UMAP visualization of 5466 single cells Control, SvX, CpG and ODN vaccinated hamsters via IM administration. Colors indicate grouping of cells into T cells, myeloid, B cells and epithelial cells based on transcriptional similarity. (**B)** Proportion of T, B, myeloid and epithelial cell types in each group. (**C)** Normalized expression of known genes on a UMAP plot to identify different cell types. **(D)** UMAP projection to visualize 17 different cell types visualized after sub clustering the major cell types at higher resolution. (**E)** Percentage of each cell subtype of T (left panel), myeloid (middle panel), B cells and epithelial cells (right panel). Significance value was calculated using ANOVA. p>0.05 was considered significant.

Seventeen cell subpopulations were distinguishable based on their expression profiles (Table S2). The immunological response to SARS-CoV-2 infection in non-vaccinated hamsters relied on innate cells such as inflammatory monocytes, neutrophils, plasmacytoid dendritic cells and natural killer T cells. In contrast, hamsters vaccinated with SvX, particularly when formulated with CpG, had a higher frequency of lymphocytes from the adaptive immune response. Specifically, both activated CD4 T cells and cytotoxic CD8 T cells highly expressing XCL1, were increased in vaccinated hamsters. Interestingly, a specific subset of B cells that does not express IgJ are significantly increased in CpG group.

The average log-fold change gene expression (avglogFC) was compared between different clusters and groups (Fig.7). In Control hamsters, genes associated with inflammation (NLRP3, IL-1B, CXCL10, CCL4, CCL8, IFI16), were one to two-fold higher in myeloid cells. In contrast, anti-inflammatory (ANXA1) and anti-viral (IFITM) genes were upregulated specifically in animals in the CpG group. Hamsters vaccinated with SvX formulated in either adjuvant, also increased the expression of CD74 conducive to B cell survival and proliferation. Without an adjuvant, however, vaccination with SvX drove the immune response towards Th2, as suggested by increased GATA-3 expression in both CD4+ and CD8+ T cells.

**Fig. 7.**
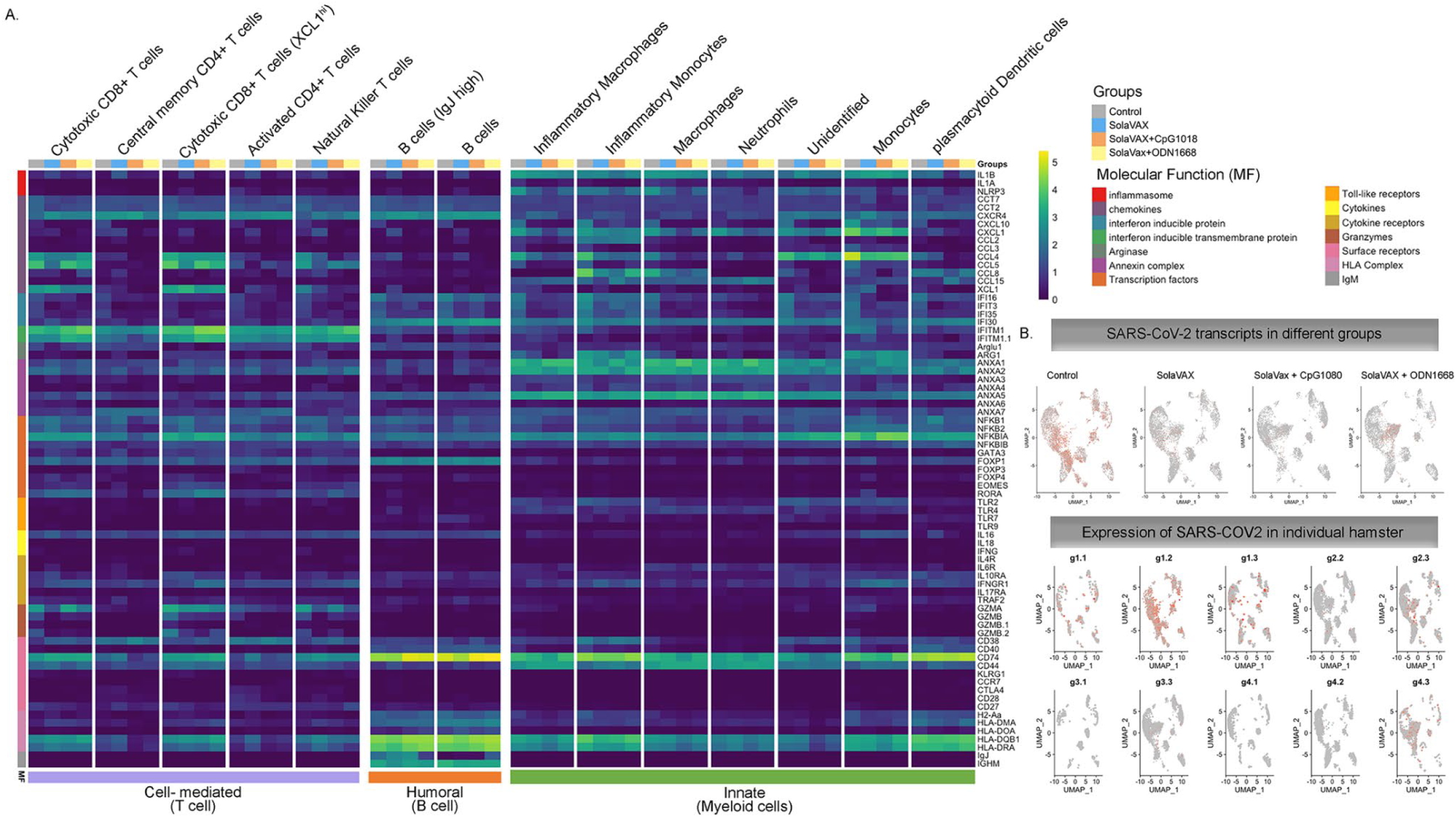
Average log fold change gene expression analysis. **(A)** Average log fold change gene expression comparing pooled gene expression between Control, SvX, CpG, and ODN for different molecular functions across different cell types. Column annotation bar at the top represents different groups, column annotation bar at the bottom represents major immune response type and row annotation bar on the left represent different molecular functions. Cell types are annotated above each cluster in the heatmap. **(B)** Expression of SARS-CoV-2 transcripts in different groups when merged (top panel) and in each individual hamster (bottom panel).

Furthermore, relevant biological functions were identified using Gene Ontology (GO) enrichment analysis of differentially expressed genes (DEGs). The top GO biological pathways were evaluated for each set of DEGs and merged within groups for p-value enrichment analysis. Pathways related to T cell differentiation, leukocyte migration, and epithelial cell development were downregulated in Control and SvX vaccinated groups but upregulated in CpG vaccinated group (Fig. 8). In contrast, viral and stress response pathways were upregulated in the Control group and the opposite trend was observed in all vaccinated groups. From a metabolic perspective, oxidative phosphorylation was suppressed in both Control and SvX hamsters; however, it was activated in the CpG group.

**Fig. 8.**
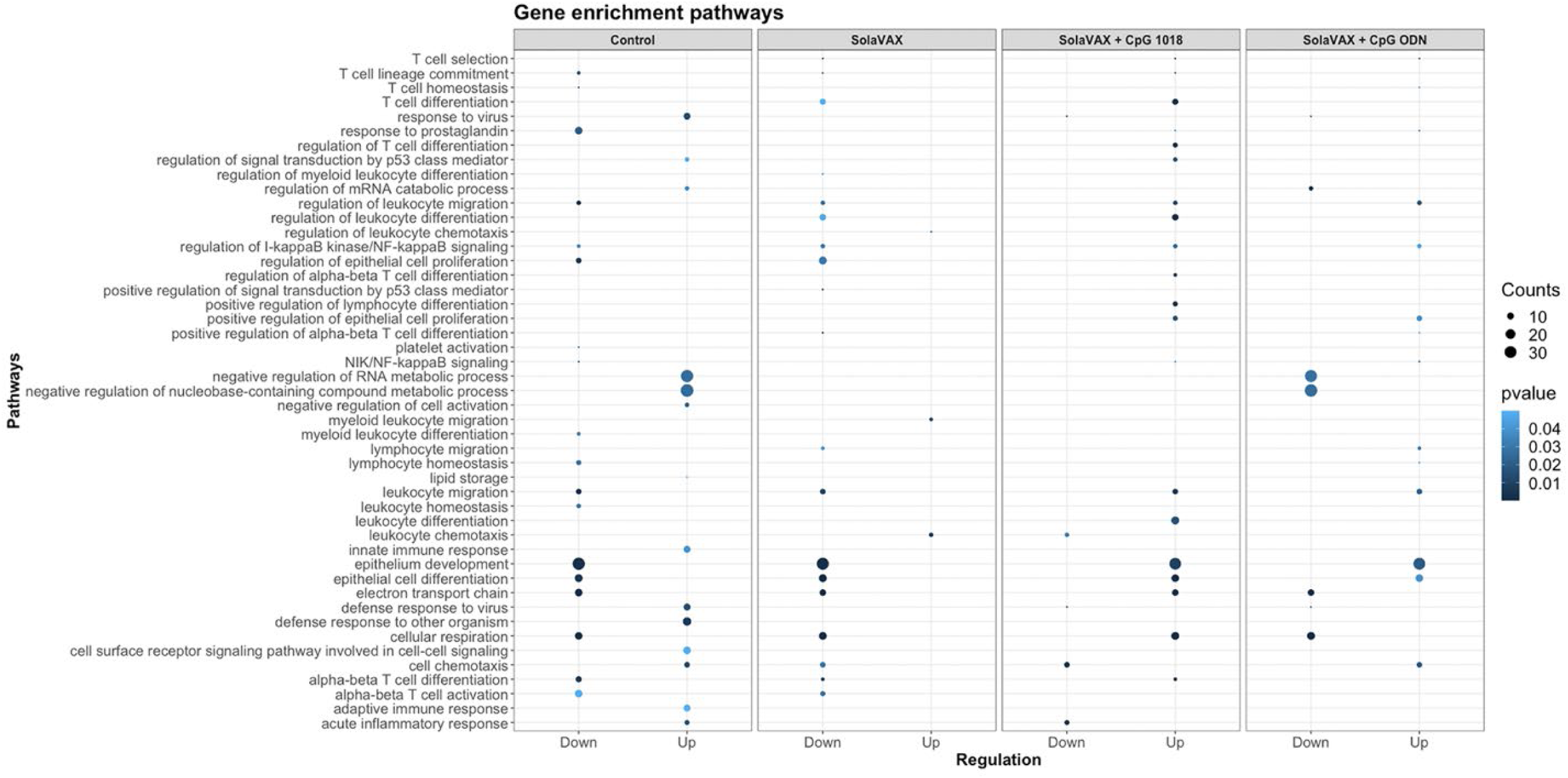
Enrichment p values for the selected Gene Ontology (GO) biological pathways of differentially expressed genes between Control and SolaVAX vaccinated hamsters, with or without adjuvant. Circles represent normalized enrichment score (NES), size of the circle represents the number of genes involved in the pathway and the color represents the significance score. p < 0.05 was considered significant.

## Discussion

Many routes to the preparation of vaccine candidates exist. All possess potential benefits and drawbacks. We evaluated the ability of a photochemical process for inactivation of pathogens in blood products to use in production of an inactivated SARS-CoV-2 whole virion for prevention of COVID-19 infection.

Our motivation for studying this approach was based on the hypothesis that the ability to inactivate virus replication without inducing damage to protein epitopes, could result in the generation of a potent vaccine candidate with intact protein antigen targets comparable to native, live-type virus. Such a candidate was hypothesized to have several advantages including the use of chemical reagents with extensive safety toxicology and general handling benefits, equipment and disposables that are currently in routine use for producing human transfusion products and a cost profile in production that could be favorable for mass production and provide for both global affordability and availability.

Our studies have demonstrated the ability of this process to inactivate SARS-CoV-2 virus via a specific, targeted guanine base modification. This work has also demonstrated the ability of products made via this method to induce a potent immune response to vaccination. This response triggered both Th1 and Th2 type immune pathways, leading to generation of neutralizing antibodies and cellular responses capable of protecting vaccinated animals against intranasal challenge with 10^5^ pfu SARS-CoV-2. The use of adjuvants was found to boost the levels of neutralizing antibody titers. Interestingly, the non-adjuvanted formulation still provided sufficient protection to prevent viral production and shedding in challenged animals. Robbianni, et al. [23] have observed varying overall neutralizing antibody levels in plasma of convalescent COVID-19 patients with consistent levels of specific sub-types against RBD epitopes. They have speculated that a subclass of antibody against receptor binding domain (RBD) epitopes may be critical in conferring therapeutic benefit in those products. The extent to which this may play a role in vaccine efficacy is unknown.

Adjuvanted formulations, particularly CpG 1018 demonstrated the lowest levels of viral shedding, preservation of normal lung morphology and airway passage integrity and reduced numbers of infiltrates in the trachea and lung tissue. Both adjuvants used in this study are known to promote Th1 immune pathway responses. Prior work with vaccine candidates suggested that ADE leading to lung immunopathology might be avoided by using Th1 promoting adjuvants [24]. Results observed here are consistent with those observations. Further studies with Th2 promoting adjuvant formulations such as alum are required, however, to further elucidate the significance of these findings.

A vaccine candidate for COVID-19 produced by this method has been demonstrated to be effective in providing protection against challenge infection in a sensitive hamster model. Importantly, we believe that such a production method could be applied to vaccine candidates targeting other viral, bacterial, and parasitic pathogens. We have already applied such an approach to the generation of solid tumor vaccines and demonstrated their use in both murine and canine disease models [24,25].

The use of this methodology may afford a means to rapidly produce vaccine candidates in response to both emergent and existing disease threats. Given the nature of the photosensitizer (riboflavin) and equipment utilized in this setting, such an approach may afford a facile method to prepare vaccine candidates in a logistically practical and cost-effective manner that avoids issues associated with current methods for production of inactivated vaccines [26]. The more selective method of nucleic acid modification without extensive protein alteration which is known to occur with current inactivation approaches may also result in more effective vaccination at lower immunogen dose, thus facilitating vaccine distribution and availability in diverse regions of the global community. These potential applications and features warrant additional testing and evaluation to fully establish their utility

## Materials and Methods

### Study Design

The objective of this study was to determine and characterize the efficacy of a novel method for creating an inactivated whole virion vaccine (SolaVAX) against SARS-CoV-2 in hamsters. Hamster group sample sizes were determined based on previous experience, using four hamsters per cohort (vaccine formulation and administration route) to evaluate the performance of the vaccine candidate in response to viral challenge using different routes of administration and in combination with different adjuvants. The sample size was large enough to demonstrate differences between treatment conditions. No animals were excluded from the analyses (1 animal death occurred in the non-adjuvanted group [SvX, SC] prior to administration of the 2^nd^ vaccine dose due to factors not related to the vaccine), and all animals were randomized to the different treatment groups. Histopathology analyses were performed blinded to the experimental cohort conditions. End points were selected prior to initiation of the study and were selected based on the objective of determining the immune responses to vaccination with the SolaVAX vaccine.

### SARS-CoV-2 virus

All virus propagation occurred in a BSL-3 laboratory setting. Virus (isolate USA-WA1/2020) was acquired through BEI Resources (product NR-52281) and amplified in Vero C1008 (Vero E6) cell culture. Vero E6 cells (ATCC CRL-1568) were cultured in Dulbecco’s modified Eagle’s medium (DMEM) supplemented with glucose, L-glutamine, sodium pyruvate, 5% fetal bovine serum (FBS) and antibiotics. Inoculation of Vero E6 cells with SARS-CoV-2 was carried out directly in DMEM containing 1% FBS. Medium harvested from infected cells 3-4 days after inoculation was clarified by centrifugation at 800 x g, supplemented with FBS to 10% and frozen to -80°C in aliquots. The virus titer was determined using a standard double overlay plaque assay.

### Viral Inactivation

Viral stock in DMEM with 10% FBS was dispensed into an illumination bag (Mirasol Illumination Bag, Terumo BCT, Lakewood, CO). Riboflavin solution (500 µmol/L) was added, residual air was removed from the product, and the bag was placed into the illuminator (Mirasol PRT System, Terumo BCT, Lakewood, CO) for treatment with UV light (150 Joules). Upon successful completion of the illumination process, the product was removed from the illuminator for vaccine preparation and characterization.

### Sequence validation of SARS-CoV-2 isolate

RNA from cell culture supernatant was extracted using the Trizol reagent (Life Technologies) according to the manufacturer’s protocol. Libraries were prepared from total RNA using the Kapa Biosystems RNA HyperPrep kit and sequenced on an Illumina NextSeq instrument to generate single end 150 base reads. Reads were mapped to the USA-WA1 reference sequence (GenBank accession MN985325.1) using bowtie2 [27]. The position, frequency, and predicted coding impact of variants were tabulated as previously described [28].

### Assessment of RNA damage following photoinactivation

Libraries were created from RNA of photoinactivated vaccine material and from matching untreated material as above. Reads were mapped to a SARS-CoV-2 reference sequence that corresponded to the consensus sequence of the virus used for vaccine production using bowtie2. The frequencies of nucleotide substitutions with basecall quality scores ≥ 30 were tabulated and normalized to the number of occurrences of the mutated bases in all reads. Analysis scripts are available at: *https://github.com/stenglein-lab/SolaVAX_sequence_analysis*.

### Vaccine concentration and preparation

Inactivated virus was concentrated using Amicon Ultra Centrifugal Filter units (Millipore Sigma) at 100k cutoff. Concentrated vaccine material was tested by plaque assay to insure complete virus inactivation. PCR was performed to determine RNA copies/mL using the Superscript III Platinum One-Step qRT-PCR system (Invitrogen). Standard curves were obtained by using a quantitative PCR (qPCR) extraction control from the original WA1/2020WY96 SARS-COV-2 isolate. Based on the ratio of pfu to virus RNA copy number for the pre-inactivation vaccine, and RNA copy number of the inactivated and concentrated vaccine, we estimated that prime and booster doses of vaccine used in hamsters were equivalent to 2.2e6 and 1.8e6 PFU-equivalents. We calculated that the amount of virus utilized per dose was on the order of 15 ng (prime) and 13 ng (boost) of virus material, respectively, in each preparation.

Vaccines were prepared immediately prior to vaccination. For the non-adjuvanted vaccine, the inactivated vaccine material was mixed equally with sterile PBS. Adjuvant CpG 1018 (Dynavax, Lot 1-FIN-3272) was mixed with equal parts of dH_2_0 then mixed with inactivated vaccine at a 1:1 ratio. One mg of adjuvant ODN1668 (Enzo ALX-746-051-M001) powder was reconstituted in 1 mL of dH_2_0 then mixed with inactivated vaccine at 1:1 ratio. Each adjuvant was utilized according to manufacturer’s recommendations for dose. For CpG 1018, 150 μg of adjuvant was used per dose. For ODN1668, 50 µg of adjuvant was used per dose.

### Animals

All hamsters were held at Colorado State University in Association for Assessment and Accreditation of Laboratory Animal Care (AAALAC) International accredited animal facilities. Animal testing and research received ethical approval by the Institutional Animal Care and Use Committee (IACUC) (protocol #18-1234A). A total of 32 male Golden Syrian hamsters (*Mesocricetus auratus)* at 6 weeks of age were acquired from Charles River Laboratories (Wilmington, MA). Hamsters were maintained in a Biosafety Level-2 (BSL-2) animal facility at the Regional Biocontainment Lab at Colorado State University during the vaccination period. The hamsters were group-housed and fed a commercial diet with access to water ad libitum. Each hamster was ear notched for animal identification. As previously described, the hamsters were randomly divided into 4 treatment groups (8 hamsters per group): Control hamsters received no vaccination, a second group (SvX) received inactivated vaccine (SolaVAX) with no adjuvant, a third group (CpG) received inactivated vaccine with CpG 1018 adjuvant, and the final group of hamsters (ODN) received inactivated vaccine with ODN1668 adjuvant. Within each group hamsters were divided in two subgroups (4 hamsters per subgroup) where one subgroup was vaccinated by subcutaneous (SC) route and the second subgroup by intramuscular (IM) route (Fig.1D).

### Clinical observations

Body weights were recorded one day before vaccination, at time of prime vaccination and booster vaccination, and then daily after challenge. Hamsters were observed daily post-vaccination for the duration of the study. Clinical evaluation included temperament, ocular discharge, nasal discharge, weight loss, coughing/sneezing, dyspnea, lethargy, anorexia, and moribund.

### Vaccination

Prior to vaccination, blood was collected from all hamsters under anesthesia and sera isolated. Each hamster in the vaccinated groups received 100 µL of vaccine (15 ng). No vaccination was administered to Control hamsters. The hamsters were maintained and monitored for 21 days. Prior to the second (booster) vaccination, blood was collected from all hamsters again under anesthesia and sera isolated. A booster vaccination (13 ng) was administered to hamsters as described in the prime vaccination. Hamsters were again maintained and monitored for additional 21 days.

### Virus challenge

All hamsters were transferred to a Biosafety Level-3 animal facility at the Regional Biocontainment Lab at Colorado State University prior to live virus challenge. Hamsters were bled under anesthesia and sera collected prior to live virus challenge to determine antibody response post vaccination.

SARS-CoV-2 virus was diluted in phosphate buffered saline (PBS) to 1 x 10^5^ pfu/mL. The hamsters were first lightly anesthetized with 10 mg of ketamine hydrochloride and 1 mg of xylazine hydrochloride. Each hamster was administered virus via 200 µL pipette into the nares (50 µL/nare) for a total volume of 100 µL per hamster. Virus back-titration was performed on Vero E6 cells immediately following inoculation. Hamsters were observed until fully recovered from anesthesia. All hamsters were maintained for three days then humanely euthanized and necropsied.

Oropharyngeal swabs were also taken prior to live virus challenge and days 1-3 after challenge to evaluate viral shedding. Swabs were placed in BA-1 medium (Tris-buffered MEM containing 1% BSA) supplemented with antibiotics then stored at -80°C until further analysis.

### Virus titration

Plaque assays were used to quantify infectious virus in oropharyngeal swabs and tissues. Briefly, all samples were serially diluted 10-fold in BA-1 media supplemented with antibiotics. Confluent Vero E6 cell monolayers were grown in 6-well tissue culture plates. The growth media was removed from the cell monolayers and washed with PBS immediately prior to inoculation. Each well was inoculated with 0.1 mL of the appropriate diluted sample. The plates were rocked every 10-15 minutes for 45 minutes and then overlaid with 0.5% agarose in media with 7.5% bicarbonate and incubated for 1 day at 37°C, 5% CO_2_. A second overlay with neutral red dye was added at 24 hours and plaques were counted at 48-72 hours post-plating. Viral titers are reported as the log_10_ pfu per swab or gram (g). Samples were considered negative for infectious virus if viral titers reached the limit of detection (LOD). The theoretical limit of detection was calculated using the following equation:

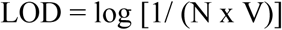

where N is the number of replicates per sample at the lowest dilution tested; V is the volume used for viral enumeration (volume inoculated/well in mL). For oropharyngeal swabs the LOD was 10 pfu/swab or 1.0 log_10_ pfu/swab. For tissues the LOD was 100 pfu/g or 2.0 log_10_ pfu/g.

### Plaque Reduction Neutralization Test

The production of neutralizing antibodies was determined by plaque reduction neutralization test. Briefly, serum was first heat-inactivated for 30 minutes at 56°C in a waterbath. Then serum samples were diluted two-fold in BA-1 media starting at a 1:5 dilution on a 96-well plate. An equal volume of SARS-CoV-2 virus (isolate USA-WA1/2020) was added to the serum dilutions and the sample-virus mixture was gently mixed. The plates were incubated for 1-hour at 37°C. Following incubation, serum-virus mixtures were plated onto Vero E6 plates as described for virus plaque assays. Antibody titers were recorded as the reciprocal of the highest dilution in which >90% of virus was neutralized. All hamsters were tested for the presence of antibodies against SARS-CoV-2 prior to vaccination.

### ELISA for anti S1, S2 and RBD antibodies

ELISA was performed to evaluate antibody binding to SARS-CoV-2 spike protein region S1 (16-685 amino acids), S2 (686-1213 amino acids), and RBD (319-541 amino acids) (all recombinant proteins from SinoBiological, Wayne, PA). The procedure was adapted from Robbiani et al. [23], with few modifications. Briefly, high binding 96-well plates (Corning, St. Louis, MO) were coated with 50 ng of S1, S2, and RBD protein prepared in PBS and incubated overnight at 4°C. Plates were washed 5 times with PBS + 0.05% Tween 20 (Sigma, St. Louis, MO) and incubated with blocking buffer (PBS + 2% BSA + 2% normal goat serum + 0.05% Tween 20) for 2 hours at room temperature (RT). Serial dilutions (1/250, 1/1250, and 1/6250) of serum obtained from naïve, non-vaccinated and vaccinated hamsters were prepared in blocking buffer and added to the plates for 1 hour. After washing, 1:10,000 dilution of HRP conjugated anti-hamster IgG (H+L) secondary antibody (Jackson Immuno Research, 107-035-142) prepared in blocking buffer was added and incubated for 1 hour. Plates were washed, TMB substrate (ThermoFisher, Waltham, MA) added, and the reaction was stopped after 10 minutes by adding 1M H_2_SO_4_. Absorbance was measured at 450 nm using a Biotek Synergy 2 plate reader (Winnoski, VT).

### Tissue Collection

Necropsies were performed three days after live virus challenge and gross pathological changes were noted. Tissues were collected for virus quantification, histopathology, flow cytometry, and single cell sequencing. For virus quantitation, portions of right cranial lung lobe, right caudal lung lobe, trachea, and nasal turbinate specimens from each hamster were weighed (100 mg per specimen) and homogenized in BA-1 media with antibiotics then frozen to -80°C until the time of analysis. The tissue homogenates were briefly centrifuged and virus titers in the clarified fluid was determined by plaque assay. Viral titers of tissue homogenates are expressed as pfu/g (log_10_). For histopathology, portions of the right cranial lung lobe, right caudal lung lobe, trachea, nasal turbinates, and spleen were collected from each hamster. The tissues were placed in 10% neutral buffered formalin for seven days then paraffin embedded and stained with hematoxylin and eosin using routine methods for histological examination. For flow cytometry and single cell sequencing, a portion of the left cranial lung lobe and spleen from each hamster was placed in PBS and immediately processed for analysis.

### Histopathology

Histopathology was blindly interpreted by a board-certified veterinary pathologist (Podell BK) at Colorado State University. The H&E slides were evaluated for morphological evidence of inflammatory-mediated pathology in lung, trachea, heart and spleen, and reduction or absence of pathological features used as an indicator of vaccine-associated protection. Each hamster was assigned a score of 0-3 based on absent, mild, moderate, or severe manifestation, respectively, for each manifestation of pulmonary pathology including mural bronchial inflammation, neutrophilic bronchitis, consolidating pneumonia, and interstitial alveolar thickening, then the sum of all scores provided for each hamster.

### Processing of lungs, spleen and blood

Lungs and spleens were processed as described by Fox et al. [29]. Briefly, a portion of the left cranial lung lobe and spleen from each hamster were aseptically transferred from PBS into DMEM then teased apart. Lungs were treated with a solution of DNase IV (500 units/mL) and Liberase (0.5 mg/mL) for 30 minutes at 37°C to dissociate and digest collagen. Both lung and spleen cells were homogenized using a syringe plunger and passed through a 70 µm filter to prepare single cell suspension. Erythrocytes were lysed using Gey’s RBC lysis buffer (0.15 M NH_4_Cl, 10 mM HCO_3_) and cells were resuspended in 1 mL of complete media.

For blood, buffy coat was harvested by adding equal volume of PBS + 2% FBS to the blood and centrifuging at 800 x g for 10 minutes at 25°C with brakes off. The buffy coat was collected and washed, and erythrocytes were lysed using 1x Miltenyi RBC lysis buffer (Miltenyi, CA). Cells were washed and resuspended in 1 mL complete media. After adding absolute counting beads (Invitrogen), total cell numbers of lung, spleen and blood were determined by flow cytometry analysis using an LSR-II (BD).

### Flow cytometry staining

Flow cytometry staining was performed as mentioned by Fox et al. [29]. Briefly, 2 x 10^6^ cells were added into each well of a 96-well v-bottom plate and incubated with 1x Brefeldin A at 37°C for 4 hours. Cells were washed and stained with Zombie NIR live/dead stain, washed and further stained with predetermined optimal titrations of specific surface antibodies (Table S3) and fluorescence minus one (FMOs). For intracellular staining, cells were further incubated with 1x Foxp3 Perm/Fix buffer (eBiosciences, San Diego, CA) for 1 hour at 37°C, washed with 1x permeabilization buffer (eBiosciences, San Diego, CA) twice and stained with intracellular antibodies cocktail (prepared in 1x permeabilization buffer) and respective FMOs overnight at 4°C. The next day, cells were washed twice and resuspended in 300 μL of 1x Permeabilization buffer. Samples were acquired using a Cytek Aurora ™ spectral flow cytometer where 100,000 events were recorded.

### Single cell RNA sequencing

Lungs cells were prepared as described above, filtered, washed and resuspended in PBS + 0.4% BSA. Cells were counted using a hemocytometer, and ∼12,000 cells were added to the 10x Genomics chromium Next GEM Chip for a target recovery of 8,000 cells. GEMs were placed in a thermal cycler and cDNA purified using Dynabeads. cDNA amplification was done using 10x Genomics single cell v3’ chemistry as per the manufacturer’s recommendations. The amplification PCR was set at 11 cycles and to eliminate any traces of primer-dimers, the PCR amplified cDNA product was purified using 0.6x SPRI beads (Beckman Coulter) before using the DNA for sequencing library preparation. Quality and quantity of cDNA was determined via Agilent TapeStation analysis using a HS-D5000 screen tape (Fig. S3). Twenty-five percent (25%) of the total cDNA amount was carried forward to generate barcoded sequencing libraries with 10 cycles of Sample Index PCR in 35-mL reaction volume (Fig. S4). Libraries were then pooled at equal molar concentration (Fig. S5) and sequenced on an Illumina NextSeq 500 sequencer to obtain a total of 941M read pairs (Illumina). An average of 78M read pairs per sample were generated with a standard deviation of 10.7M read pairs. Low-quality cells with <200 genes/cell and cells that express mitochondrial genes in >15% of their total gene expression were excluded. Gene expression in each group was normalized based on the total read count and log transformed.

Sequenced samples were de-multiplexed using Cell Ranger mkfastq (Cell Ranger 10× Genomics, v3.0.2) to generate fastq files and aligned to the *Mesocricetus auratus* (accession GCA_000349665) and SARS-CoV-2 (reference genome MN985325) reference genomes using CellRanger count pipeline. Filtered barcode matrices were analyzed by Seurat package Version 3.0. Low quality cells, defined as expressing <200 genes/cell or those in which mitochondrial genes corresponded to >15% of their total gene expression, were excluded. Samples within groups were merged and downsampled to the same number of cells per group. Thereafter, gene expression for each group was normalized based on total read counts and log transformed. All groups were integrated using Seurat integration strategy [30], aligned samples scaled, and cells analyzed by unsupervised clustering (0.5 resolution), after principal components analysis (PCA). The top 15 principal components were visualized using UMAP. Differentially up-regulated genes in each cluster were selected with >0.25 log fold change and an adjusted p < 0.05. Cell types were assigned by manually inspecting the top 20 upregulated genes, in addition to identifying previously published specific markers such as FSCN1 and GZMA for dendritic cells and CD8+ effector T cells, respectively [31]. Differentially expressed genes (DEGs) between non-vaccinated group and vaccinated groups were identified using DESeq2 algorithm, with a Bonferroni-adjusted p< 0.05 and a log2 fold change > 1.

### Statistical Analyses

Group mean (n=4) viral titers were analyzed using a two-way ANOVA analysis followed by a post hoc test to analyze differences between the Control group and the vaccinated groups. In the case where samples reached the LOD, values were entered as 0 for statistical analysis. Data were considered significant if *p*<0.05. Analysis was performed using GraphPad Prism software (version 8.4.2) (GraphPad Software, Inc, La Joia, CA). Mean (n=4) subjective pathology scores were compared between groups using the Kruskall-Wallis test for non-parametric data with an alpha of 0.05. The flow cytometry results were analyzed using FlowJo as well as a newly published methodology [29]. Events were filtered to cell populations that constituted greater than 2% of the live leukocytes of at least one sample, where a cell population is defined by the combination of positive and negative markers. Statistical significance among A and among B groups was determined using Anova and Tukey Honest Significant Difference.

## Acknowledgments

The authors would like to thank the Flow Cytometry, Next Generation Sequencing, and Experimental Pathology Core laboratories at Colorado State University for their assistance and expertise with sample analyses. Additionally, the authors thank Ann Hess and the Graybill Statistical Laboratory at Colorado State University for assistance with statistical analyses of the viral titer and PRNT data. We thank BEI for providing the SARS-CoV-2 virus utilized in these studies. The reagent was deposited by the Centers for Disease Control and Prevention and obtained through BEI Resources, NIAID, NIH: SARS-Related Coronavirus 2, Isolate USA-WA1/2020, NR-52281. We thank Dynavax for providing the CpG 1018 adjuvant. Vero African Green Monkey Kidney Cells (ATCC® CCL-81™), FR-243, were obtained through the International Reagent Resource, Influenza Division, WHO Collaborating Center for Surveillance, Epidemiology and Control of Influenza, Centers for Disease Control and Prevention, Atlanta, GA, USA. The RNA extraction control was obtained through BEI Resources, NIAID, NIH: Quantitative PCR (qPCR) Extraction Control from Inactivated SARS Coronavirus, Urbani, NR-52349.

## Data and materials availability

All data associated with this study are present in the paper or the Supplementary Materials. SARS-CoV-2 USA-WA1 strain sequencing data may be accessed via GenBank accession MN985325.1. Analysis scripts for RNA damage following photoinactivation are available at: *https://github.com/stenglein-lab/SolaVAX_sequence_analysis*.

## Supplementary Materials

**Fig. S1. Semiquantitative lung pathology scores from all study groups separated by route of administration.**

**Fig. S2. Statistically non-significant flow cytometry populations within intramuscular and subcutaneously vaccinated groups.**

**Fig. S3. cDNA amplification traces**

**Fig. S4. CDNA library traces**

**Fig S5. Total cDNA concentration and cDNA library molarity of individual samples.**

**Table S1. Markers for Identification of different cell types**

**Table S2. Consensus-changing mutations in the SARS-CoV-2 isolate used for hamster infection studies relative to the USA-WA1 sequence**

**Table S3: Flow cytometry panel to Th1 and Th2 response**

**Table S1.** Consensus-changing mutations in the SARS-CoV-2 isolate used for hamster infection studies relative to the USA-WA1 900 sequence (accession MN985325.1).

| **Position in genome**<br>**(nt)<sup>1</sup>** | **Gene** | **Nucleotide**<br>**substitution** | **Amino acid**<br>**substitution** | **Variant frequency<sup>2</sup>** |
| --- | --- | --- | --- | --- |
| 13845 | nsp12 | U → G | D135E | 0.86 |
| 22205 | S | G → C | D215H | 0.90 |
| 23616 | S | G → A | R685H | 0.87 |
| 26542 | M | C → U | T7I | 0.93 |
| 28853 | N | U → A | S194T | 0.94 |

1. Coordinates relative to MN985325.1
2. Fraction of reads with alternate base

**Fig. S1.**
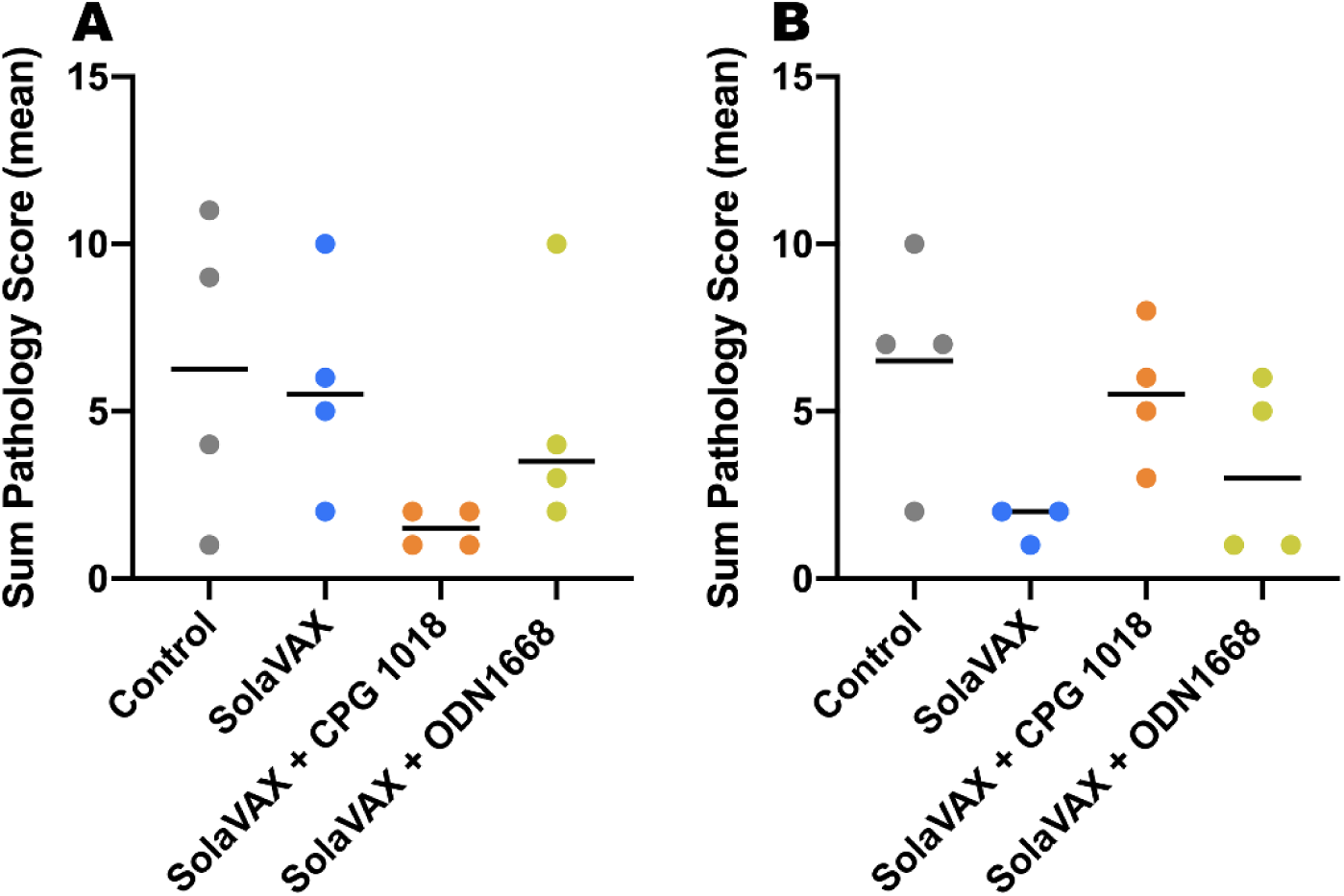
Semiquantitative lung pathology scores from all study groups separated by route of administration. Overall severity of lung pathology was determined by the sum of severity scores for four pathological features with 12 being the maximum assigned sum of severity scores. Data are shown for all groups, separated by intramuscular **(A)** and subcutaneous **(B)** routes of immunization. Data points represent sum scores of individual animals with the bar representing the mean.

**Fig. S2.**
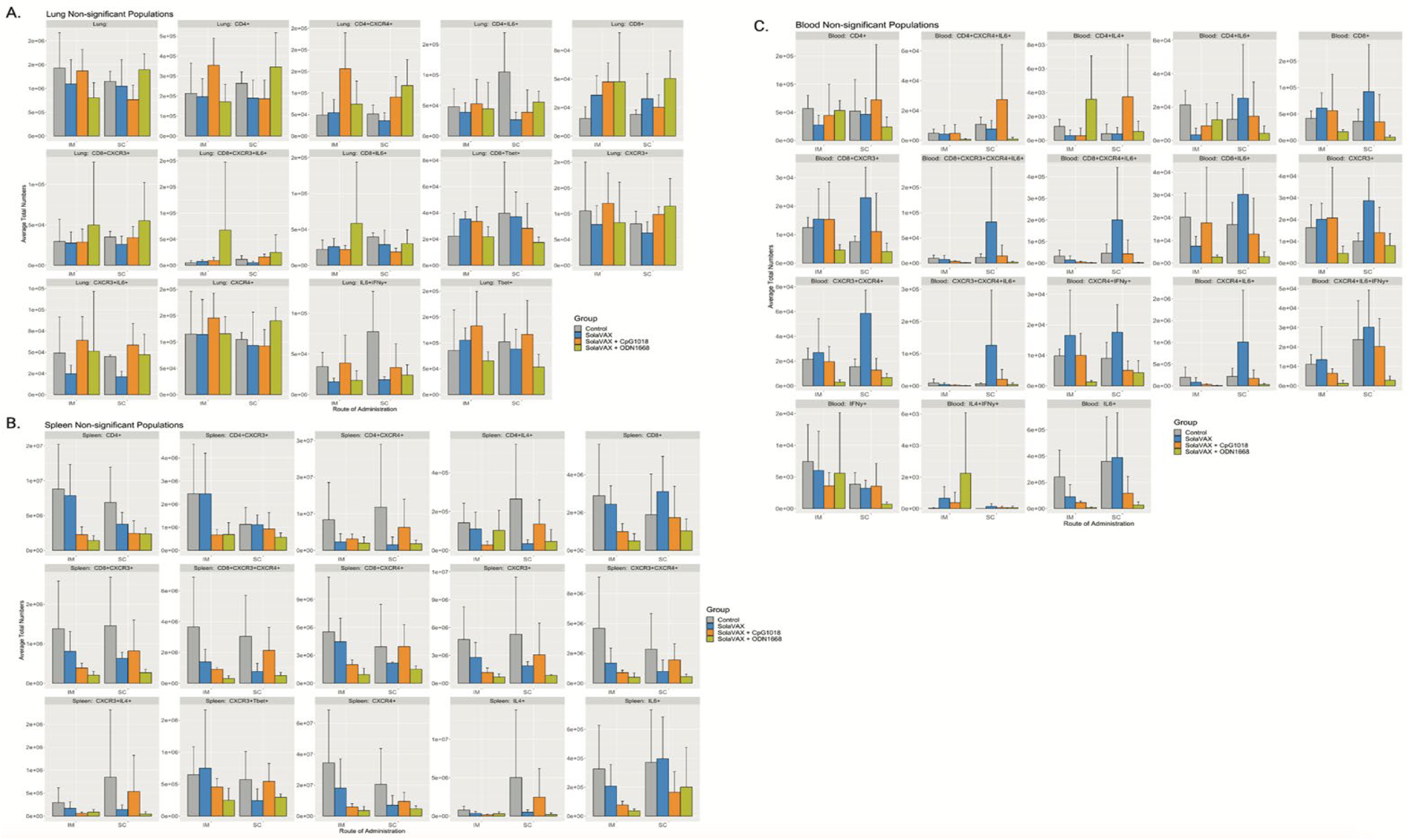
Statistically non-significant flow cytometry populations within intramuscular and subcutaneously vaccinated groups. The bar plots show the statistically non-significant populations for the lung **(A)**, spleen **(B)**, and blood **(C)**. The y-axis displays the average total numbers of cells for the eight groups. The population names at the top of the plots indicate the positive markers in the population: the population is negative for all other markers in the panel.

**Table S2.** Markers for Identification of different cell types. By using principal component analysis, 17 clusters were generated by using Seurat pipeline. These clusters were classified into different cell types based on specific markers cited in literatures.

| Markers | Cell type | References |
| --- | --- | --- |
| Marco, CD86, CD274, NLRP3, IL-1B | Inflammatory Macrophages | [30] |
| CD14, Saa3, THBS1, CCL8 | Inflammatory monocytes | [30, 31] |
| Marco, CD80, FABP5 | Macrophages | [30] |
| ELANE, NET1, S100A6 | Neutrophils | [32] |
| S100A8, S100A9, CD14 | Monocytes | [31] |
| TCF4, FSCN1, CD83 | plasmacytoid dendritic cells | [33, 34] |
| CD3D, CD8A, GZMA, GZMB, NKG7, CD44 | Cytotoxic CD8 T cells | [33-35] |
| CD3D, CD8A, GZMA, NKG7, CD44, XCL | Cytotoxic CD8 T cells, XCL<sup>hi</sup> | [33-35] |
| CD3D, CD4, CD44, CD62L, CD38 | Central memory CD4+ T cells | [33, 34] |
| CD3D, CD4, CD44, TNFRSF4 | Activated CD4+ T cells | [33] |
| CD3D, GZMA, GZMK, *TNFRSF* | NK cells | [33, 35] |
| CD79B, CD74, H2-Aa, IGJ, IGHM | B cells IGJ<sup>hi</sup> | [33, 36] |
| CD79B, CD74, H2-Aa | B cells | [33, 36] |
| SFTPC, SFTPB | Type –II alveolar cells | [33] |
| SFTPC, SOX4 | Epithelial progenitor cells | [33] |

**Fig. S3.**
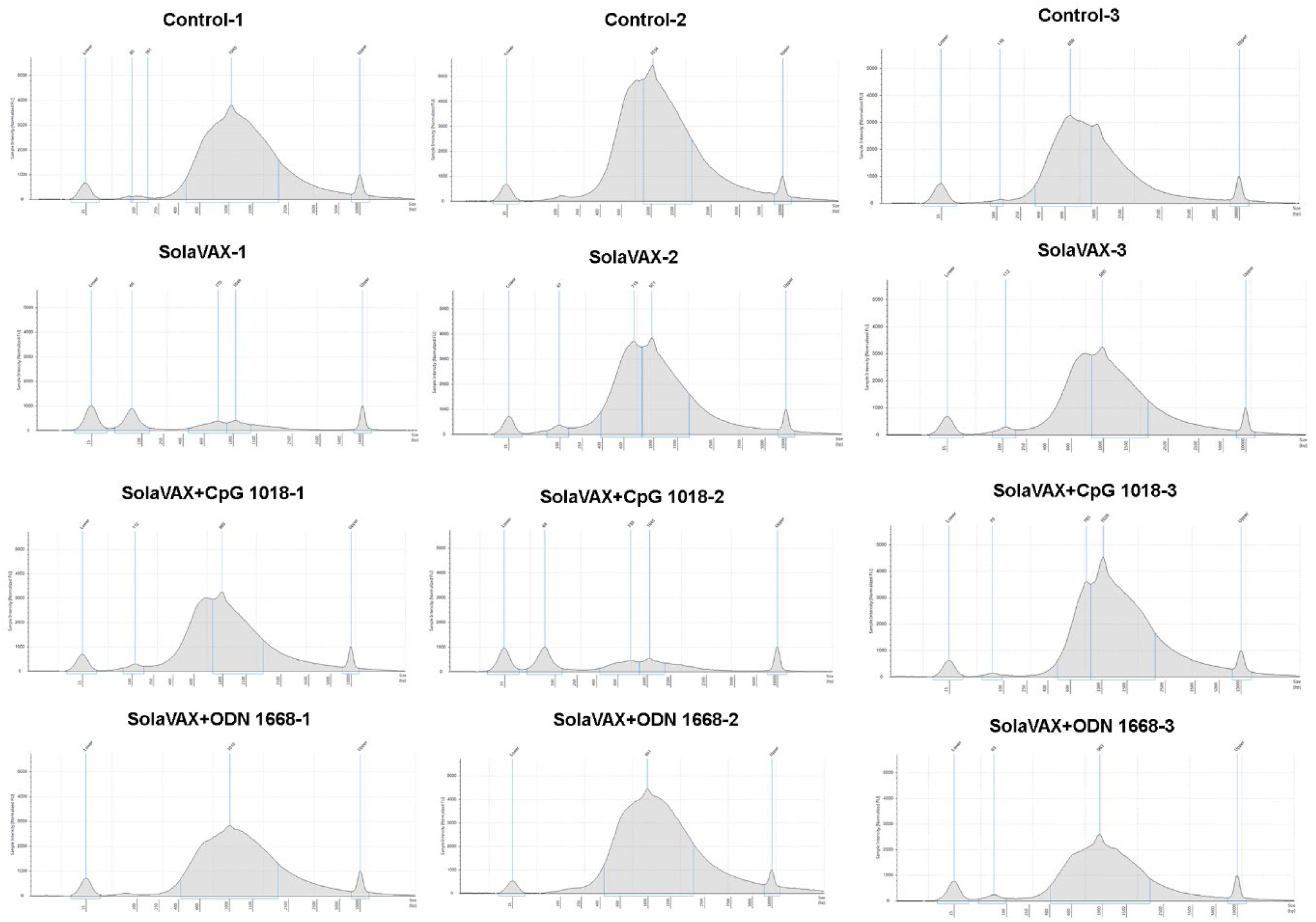
cDNA amplification traces. cDNA was amplified and quality and quantity were evaluated via Agilent Tapestation using HS-D5000 screen tapes and reagents. Traces here represent amplified cDNA after 10-fold dilution.

**Fig. S4.**
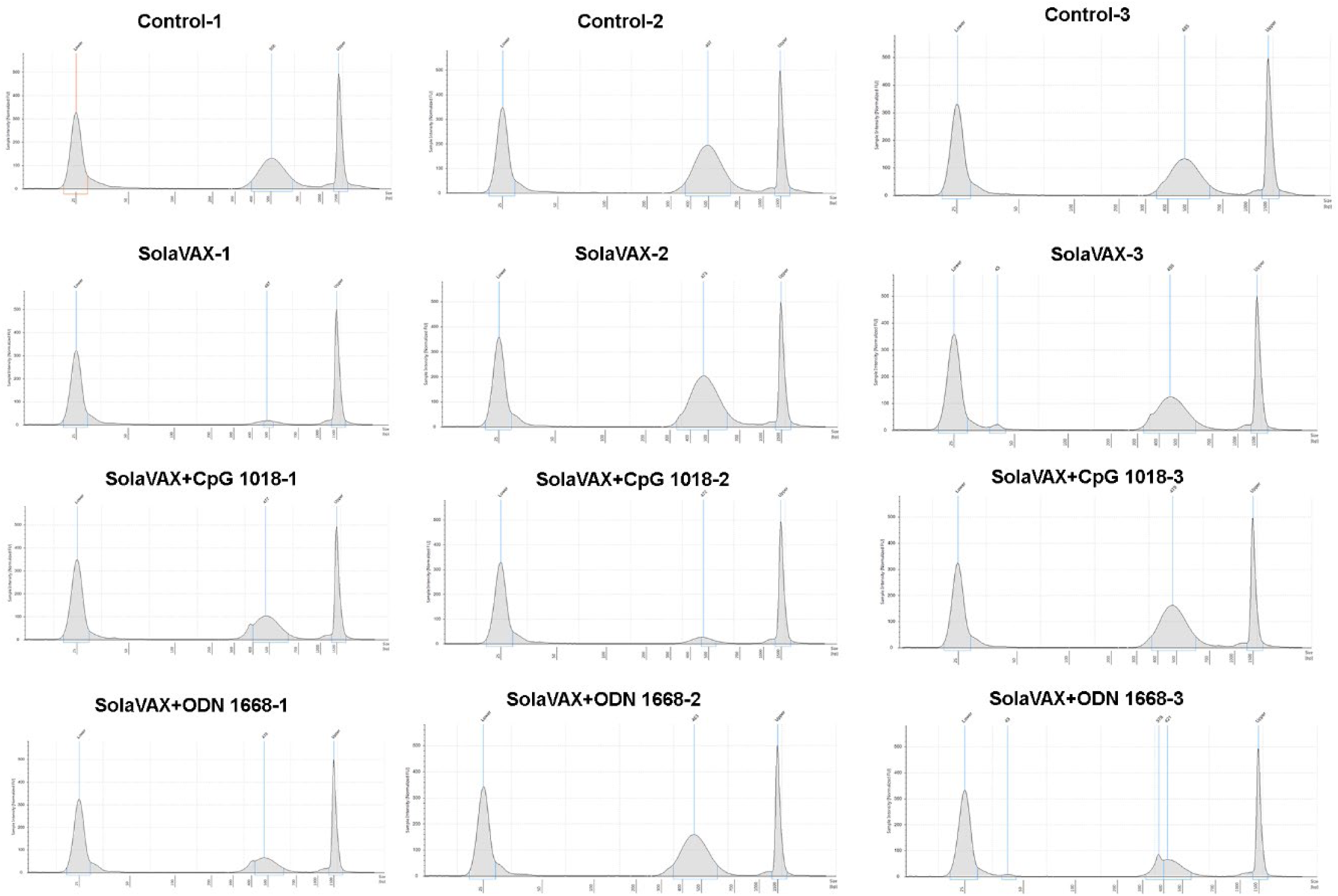
cDNA library traces. cDNA was amplified, library was prepared, and quality and quantity were evaluated via Agilent Tapestation using HS-D1000 screen tapes and reagents. Traces here represent cDNA library after 10-fold dilution.

**Fig. S5.**
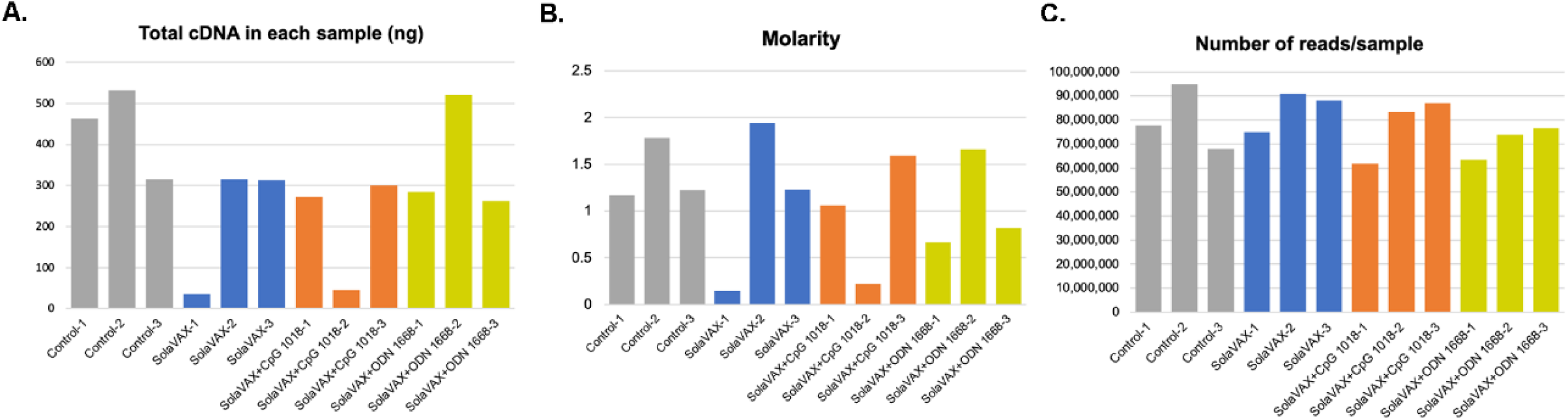
Total cDNA concentration, cDNA library molarity and number of reads for individual samples. Total cDNA in the sample was calculated by taking concentration of cDNA obtained (in pg/µL) between 200-9000 bp. Molarity of the library was evaluated using region between 250 and 1000 bp. Number of reads were obtained from the combined sequencing run performed in Illumina Next Seq 500.

**Table S3.** Flow cytometry panel: Flow cytometry panel to study the Th1 and Th2 immune responses in non-vaccinated and SolaVAX vaccinated hamsters. Antibodies were selected based on previous studies and by percent similarity between hamsters to either mouse/rat based on availability of the antibodies.

| Antibody | Clone | fluorophore | Concentration of antibody used | Company | Catalogue | References |
| --- | --- | --- | --- | --- | --- | --- |
| CD4 | GK1.5 | Pacific blue | 1 µg/mL | Biolegend | 100428 | [37] |
| CD8 | 341 | FITC | 2 µg/mL | BD Biosciences | 554973 | [38, 39] |
| IFN-γ | XMG1.2 | BV785 | 1 µg/mL | Biolegend | 505838 | [40] |
| IL-10 | JES5-16E3 | BV421 | 1 µg/mL | Biolegend | 505022 | [40] |
| Gata-3 | 16E10A23 | PE | 0.5 µg/mL | Biolegend | 653804 | [41] |
| Tbet | 4B10 | BV711 | 0.5 µg/mL | Biolegend | 644820 | [41] |
| IL-4 | 11-B11 | APC | 1 µg/mL | Biolegend | 504106 | [40] |
| TNFα | MP6-XT22 | PE-Dazzle 594 | 0.5 µg/mL | Biolegend | 506346 | Based on percent similarity of protein sequence, hamster and mouse: 82% |
| IL-6 | MP5-20F3 | Percp efluor 710 | 1 µg/mL | Thermo Scientific | 46-7061-82 | Based on percent similarity of protein sequence, hamster and mouse: 73% |
| CXCR3 | 173 | APC-Fire 750 | 1 µg/mL | Biolegend | 126540 | Based on percent similarity of protein sequence, hamster and mouse: 91.1% |
| CXCR4 | 2B11 | BV650 | 2 µg/mL | BD Biosciences | 740526 | Based on percent similarity of protein sequence, hamster and mouse: 82% |
| Zombie NIR |  | Live/dead | 1:2000 dilution | Biolegend | 423106 | Based on percent similarity of protein sequence, hamster and mouse: 93.2% |

